# Exportin Crm1 is important for Swi6 nucleocytoplasmic shuttling and MBF transcription activation in *Saccharomyces cerevisiae*

**DOI:** 10.1101/2020.06.17.155598

**Authors:** Kenneth D. Belanger, William T. Yewdell, Matthew F. Barber, Amy N. Russo, Mark A. Pettit, Emily K. Damuth, Naveen Hussain, Susan J. Geier, Karyn G. Belanger

**Author notes:** Department of Biology, University of Oregon, Eugene, OR, USA. The Estée Lauder Companies, Inc., Mellville, NY, USA. Department of Emergency Medicine, Rochester General Hospital, Rochester, NY, USA. Department of Emergency Medicine, Cooper University Health Care, Camden, NJ, USA. Department of Chemistry, Colgate University, Hamilton, NY, USA. Center for Learning, Teaching, and Research, Colgate University, Hamilton, NY, USA. Corresponding author: (KDB).

## Abstract

The Swi6 protein acts as a transcription factor in budding yeast, functioning in two different heterodimeric complexes, SBF and MBF, that activate the expression of distinct but overlapping sets of genes. Swi6 undergoes regulated changes in nucleocytoplasmic localization throughout the cell cycle that correlate with changes in gene expression. While the process of Swi6 nuclear import is well understood, mechanisms underlying its nuclear export remain unclear. Here we investigate Swi6 nuclear export and its impact on Swi6 function. We show that the exportin Crm1, in addition to three other karyopherins previously shown to affect Swi6 localization, is important for Swi6 nuclear export and activity. A truncation of Swi6 that removes a putative Crm1 nuclear export signal results in the loss of changes in nucleocytoplasmic Swi6 localization that normally occur during progression through the cell cycle. Mutagenesis of the NES-like sequence or removal of Crm1 activity using leptomycin B results in a similar decrease in nuclear export as cells enter S-phase. Using two-hybrid analysis, we also show that Swi6 associates with Crm1 *in vivo*. Alteration of the Crm1 NES in Swi6 results in a decrease in MBF-mediated gene expression, but does not affect expression of an SBF reporter, suggesting that export of Swi6 by Crm1 regulates a subset of Swi6 transcription activation activity. Finally, alteration of the Crm1 NES in Swi6 results in cells that are larger than wild type, but not to the extent of those with a complete Swi6 deletion. Expressing a Swi6 NES mutant in combination with a deletion of Msn5, an exportin involved in Swi6 nuclear export and specifically affecting SBF activation, further increases the large cell phenotype, but still not to the extent observed in a Swi6 deletion mutant. These data suggest that Swi6 has at least two different exportins, Crm1 and Msn5, each of which interacts with a distinct nuclear export signal and influences expression of a different subset of Swi6-controlled genes.

**Summary Statement:** Precise intracellular localization is important for the proper activity of proteins. Here we provide evidence that the Swi6 transcription factor important for cell cycle progression shuttles between the cell nucleus and cytoplasm, its nuclear export is important for its activity, and that it contains a nuclear export signal (NES) recognized by the Crm1 nuclear transport factor.

## Introduction

Cell cycle progression is a highly regulated process in all cells. In eukaryotes, cells become committed to progression through the cell cycle during the late G_1_ phase at a point termed the restriction point, also referred to as START in yeast (Fisher, 2016; Johnson & Skotheim, 2013). In *Saccharomyces cerevisiae*, progression through START correlates with specific cellular characteristics such as appropriate cell size and rate of protein synthesis, as well as the transcription of a certain subset of genes, including about 200 of which undergo a peak in expression at this point (Cho et al., 1998; Costanzo et al., 2003; Ferrezuelo et al., 2010; Spellman et al., 1998; Wijnen et al., 2002). These genes function in a variety of roles, including mating type switching, DNA replication, budding, cell wall synthesis, and spindle pole body duplication (see (Futcher, 2002; Jorgensen & Tyers, 2004; Stillman, 2013)). All of these molecular events are critical to proper progression through the cell cycle and depend on specific transcriptional activation at a precise stage of the cycle (Flick & Wittenberg, 2005; Futcher, 2002). This activation is initiated by the Cln3/Cdc28 cyclin/cyclin-dependent kinase (Cdk) complex, which stimulate an intricately timed sequence of protein modifications and gene expression changes that commit the cell to START (Dončić et al., 2011; Eser et al., 2011).

Of the hundreds of yeast genes that undergo changes in expression during the G_1_-S transition, the vast majority fall at least partially under the regulation of two heterodimeric transcription factors, MBF and SBF (Bean et al., 2005). MBF (<u>M</u>luI cell-cycle <u>b</u>ox binding factor) and SBF (<u>S</u>wi4/6 cell-cycle <u>b</u>ox binding factor) are both composed of Swi6 plus one other protein, Mbp1 and Swi4, respectively ((Breeden & Nasmyth, 1987; Koch et al., 1993), reviewed in (Hendler et al., 2018)). In their respective complexes with Swi6, Swi4 and Mbp1 function as the sequencespecific DNA binding component, while Swi6 itself functions as the transcription activating component (Andrews & Moore, 1992; Koch et al., 1993; Lowndes et al., 1992). SBF and MBF regulate the expression of distinct but overlapping sets of genes with MBF primarily controlling genes involved in DNA replication and repair while SBF upregulates genes involved in cell growth, cell wall organization, and the timing of progression through START ((Bean et al., 2005; de Bruin et al., 2006); reviewed in (Wittenberg & Reed, 2005)).

In eukaryotic cells, progression through the cell cycle is dependent on the precise temporal control of distinct transcriptional programs. One mechanism that contributes to this control is the regulated nucleocytoplasmic localization of specific proteins. Proteins entering the nucleus either contain a nuclear localization signal (NLS) within their polypeptide sequence or associate with an NLS-containing protein (reviewed in (Lange et al., 2007; Soniat & Chook, 2015)). The NLS is then recognized by an importin member of the Ran-binding karyopherin protein family of nuclear transport factors (NTFs), which chaperones the NLS-containing proteins through the nuclear pore complexes (NPCs) that perforate the nuclear envelope (Nima Mosammaparast & Pemberton, 2004; Xu et al., 2010). Once in the nucleus, the importin releases its cargo protein so that the cargo is free to perform its intranuclear function (Görlich et al., 2003; Lee et al., 2005). Many proteins imported into the nucleus also undergo nuclear export (Kirli et al., 2015; Mackmull et al., 2017; Thakar et al., 2013), resulting in the regulated nucleocytoplasmic shuttling of these polypeptides. Actively exported proteins contain a nuclear export signal (NES) that is recognized by an exportin member of the karyopherin NTF family, which associates with the NES and transports the cargo protein out of the nucleus through the NPCs (Fornerod et al., 1997; Kutay & Güttinger, 2005). Importantly, there are multiple types of karyopherin proteins (14 in yeast; approximately 20 in mammals; (Quan et al., 2008)) and different karyopherins function as importins or exportins for specific subsets of proteins based on the NLSs and NESs present within those proteins, thus allowing for regulation of nucleocytoplasmic localization of specific proteins or protein families.

The activity of many transcription factors is regulated in part by control of their intracellular localization. In mammalian cells, the transcriptional activation activity of NFκB is largely determined by the cytosolic regulator IκB which, when present, binds NFκB in a manner that blocks its NLS and thus prevents it from entering the nucleus and activating transcription. Upon degradation of IκB after signal-activated phosphorylation, the NFκB NLS is exposed, associates with its importin protein, and rapidly enters the nucleus (Ghosh & Karin, 2002). This nuclear entry results in activation of transcription of NFκB responsive genes (Hayden & Ghosh, 2008). Nucleocytoplasmic localization regulates a variety of other important transcription factors, including SMAD, β-catenin, p53, and STAT proteins (Derynck et al., 1998; Logan & Nusse, 2004; Meyer et al., 2002; Roth et al., 1998).

The yeast Swi6 transcription factor undergoes dynamic changes in intracellular localization, as it is primarily nuclear under some conditions and cytoplasmic under others (Kim et al., 2010; Sidorova et al., 1995; Taba et al., 1991). Two different importins, the Kap60/Kap95 (Importin-α/β) heterodimer and the monomeric Kap120 β-importin, bind to distinct NLS sequences within Swi6 to mediate its nuclear import under different conditions (Harreman et al., 2004; Kim et al., 2010). Swi6 rapidly accumulates in the nucleus in late mitosis and remains primarily nuclear until the end of G1, at which time it becomes cytosolic (Sidorova et al., 1995). The import during G1 occurs via the Kap60/Kap95 importin-α/β complex and is inhibited by Clb6/Cdc28-mediated phosphorylation of Ser160 adjacent to the classical NLS (cNLS) recognized by Kap60/Kap95 (Geymonat et al., 2004; Harreman et al., 2004). Swi6 nuclear import also rapidly occurs in response to cell wall stress, in this case being mediated by Kap120 and negatively regulated by Mpk1-mediated phosphorylation of Ser238 adjacent to the Kap120-recognized NLS (Kim et al., 2010). In both cases, phosphorylation correlates with a rapid loss of nuclear Swi6 and changes in Swi6-mediated gene expression (Geymonat et al., 2004; Harreman et al., 2004; Kim et al., 2010; Sidorova et al., 1995).

Regulation of Swi6 nuclear export also appears to be important for determining its transcriptional activity. A deletion of Msn5, a yeast karyopherin with exportin function, results in Swi6 remaining localized to the nucleus throughout the entire cell cycle, and leads to decreased expression of SBF-responsive but not MBF-responsive genes (Queralt & Igual, 2003). Interestingly, depletion of Msn5 during one round of the cell cycle results in a lack of Swi6 binding to an SBF-responsive promoter in the following G_1_-S phase, suggesting that the nuclear export of Swi6 is necessary for proper SBF function in the succeeding cell cycle (Queralt & Igual, 2003). Thus, regulation of both nuclear import and export is critical for proper Swi6-mediated gene expression.

Here we provide evidence for the function of an additional exportin, Crm1/Xpo1, in regulating the intracellular localization of Swi6 in *Saccharomyces cerevisiae* in a cell cycle dependent pattern and identify a Crm1 nuclear export signal (NES) within Swi6. We also provide evidence that both Msn5 and Crm1/Xpo1 are critical for normal Swi6 function, and that regulated export by each differentially impacts Swi6 transcription activation activity.

## Results

### Swi6-GFP undergoes nucleocytoplasmic shuttling

Swi6 protein is localized primarily in the nucleus during G1 and S-phase of the cell cycle, when it functions as an active transcription factor (Bean et al., 2005; Sidorova et al., 1995). However, this nuclear accumulation decreases during G2 and M-phase and the protein becomes largely cytoplasmic (Sidorova et al., 1995). While these observations suggest that Swi6 protein shuttles in and out of the nucleus in a cell cycle-dependent manner, such changes in localization could also be observed due to changes in nuclear import rates combined with localized Swi6 protein degradation. In order to confirm that Swi6-GFP shuttles in and out of the nucleus, we performed a heterokaryon shuttling assay (Feng & Hopper, 2002). Galactose-dependent expression of Swi6-GFP was induced in a haploid yeast strain until Swi6-GFP was observed in the nucleus, and then repressed by incubating in glucose. The cells were then mated with a yeast strain containing a *kar1-1* allele that prevents karyogamy after zygote formation (Conde & Fink, 1976). The resulting heterokaryon has Swi6-GFP in one nucleus and lacks fluorescence in the other at the time of cell fusion. Because new Swi6-GFP expression is repressed, the only way for the *kar1-1* mutant nucleus to accumulate significant fluorescence is via Swi6-GFP protein export or diffusion from the wild-type nucleus and re-import into the *kar1-1* nucleus. Thus, if only one nucleus exhibits fluorescence, Swi6-GFP does not shuttle. In contrast, if GFP is present in both nuclei, Swi6 is capable of nucleocytoplasmic shuttling. We observe that in every heterokaryon generated (n = 20), Swi6-GFP is found in both nuclei (Fig. 1). As controls for this assay, we observe that the shuttling tRNA export factor Cca1-GFP (Feng & Hopper, 2002) is present in both nuclei in heterokaryons, while the non-shuttling histone H2B protein (N. Mosammaparast et al., 2001) is retained only within one nucleus. These data indicate that Swi6-GFP truly is a “shuttling” protein that undergoes both nuclear import and export under steady-state conditions.

**Figure 1.**
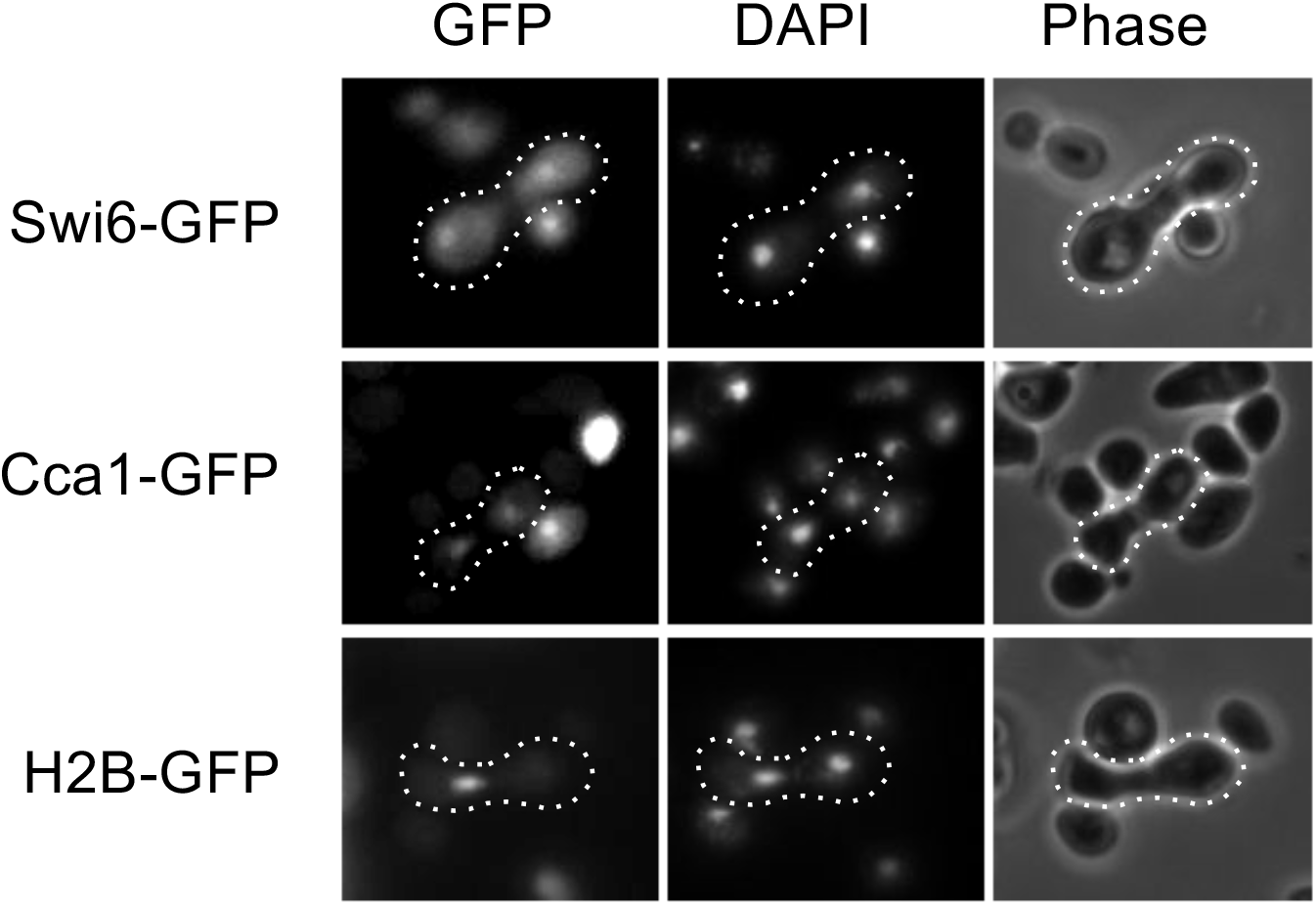
Swi6 shuttles between the nucleus and cytoplasm in *S. cerevisiae*. To test for nucleocytoplasmic shuttling, chimeric Swi6-GFP, Cca1-GFP, or histone H2B-GFP proteins were expressed under control of a *GAL1* promoter. Cells were incubated in the presence of galactose to induce expression, then incubated with glucose for 1 hr to repress expression. Cells were then mated with *kar1-1* mutant cells (MS739, (Vallen et al., 1992)) to form heterokaryons and GFP localization observed using fluorescence microscopy (left panels). Nuclear location was determined by DAPI (middle column). Cells expressing Swi6-GFP (top row), the shuttling protein Cca1-GFP (middle row), and non-shuttling histone-H2B-GFP (bottom row) are depicted, with representative zygotes highlighted by a dashed outline.

### Swi6 residues 250-258 are important for optimal nuclear export

While the redistribution of Swi6 between the nucleus and cytosol has been identified previously as necessary for changes in Swi6 activity throughout the cell cycle and multiple NLSs and importins have been identified that contribute to nuclear import (Harreman et al., 2004; Kim et al., 2010; Queralt & Igual, 2003; Sidorova et al., 1995), the region(s) of Swi6 necessary for nuclear export remain unknown. To identify sequences important for Swi6 export, we generated Swi6 truncation mutants fused with GFP, expressed each in wild type yeast, and performed fluorescence microscopy on synchronously growing populations of cells (Fig. 2). Both full-length (Swi6^1-803^; (Sidorova et al., 1995)) and truncated (Swi6^1-273^-GFP; Figs 2B and 2F) exhibit reduced nuclear localization shortly after release from a-factor, with nearly completely cytoplasmic fluorescence 80 min after release, and return to the nucleus by 120 min. In contrast, the more severely truncated version (Swi6^1-181^-GFP) is retained in the nucleus throughout the cell cycle (Figs. 2B and 2F). Thus, the region of Swi6 between amino acids 181 and 273 is important for changes in nuclear Swi6 concentration during G1/S.

**Figure 2.**
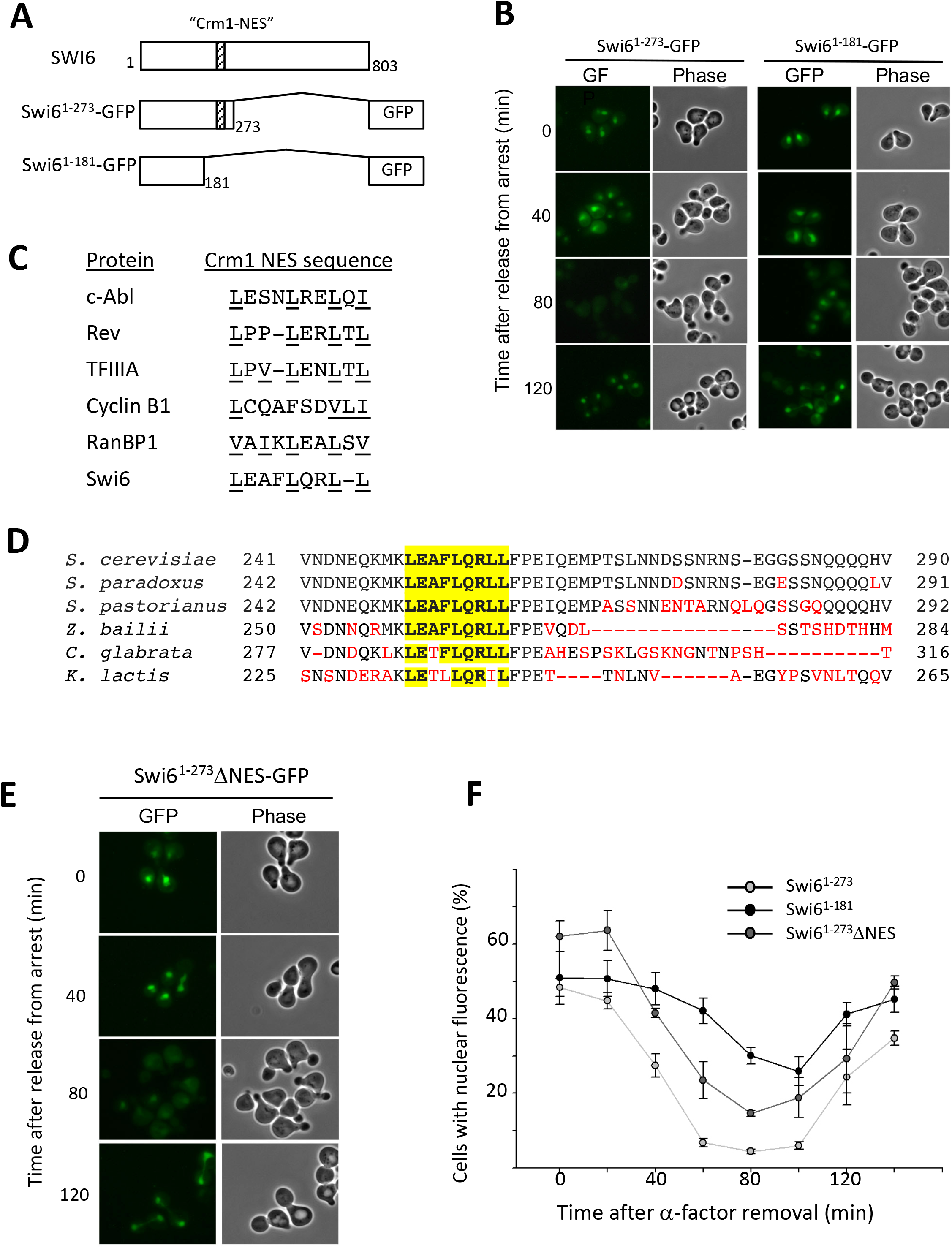
Swi6 contains a leucine-rich domain that is necessary for efficient nuclear export. (A) Cartoon diagram depicting truncation mutants lacking amino acids 274 – 803 (Swi6^1-273^-GFP) and 182 – 803 (Swi6^1-181^-GFP) from the carboxy-terminus of Swi6 fused with GFP. (B) Deletion of amino acids 182 – 274 results in decreased Swi6 nuclear export. Cells expressing Swi6^1-273^-GFP (left) or Swi6^1-181^-GFP (right) were arrested with a-factor, then released from arrest to observe Swi6 localization in synchronous cell populations. Cells expressing the Swi6-GFP fusions were observed by direct fluorescence (GFP) and phase-contrast (Phase) microscopy at the timepoints indicated after release from arrest. (C) Alignment of Swi6 amino acids 250 – 258 with leucine-rich nuclear export signals (NESs) from other shuttling proteins. Leucine residues are underlined. (D) Alignment of *Saccharomyces cerevisiae* Swi6 leucine-rich sequence and flanking residues with homologous regions from fungi *Saccharomyces paradoxus, Saccharomyces pastorianus, Zygosaccharomyces bailii, Candida glabrata*, and *Kluyveromyces lactis*. Leucine-rich Crm1-NES-like sequences aligning with amino acids 250 – 258 from *S. cerevisiae* are highlighted in yellow. Amino acids that differ from *S. cerevisiae* Swi6 are red. (E) Mutagenesis of leucines within amino acids 250 – 258 to alanines results in decreased Swi6 nuclear export during the cell cycle. A plasmid encoding Swi6 amino acids 1 – 273 fused with GFP was subjected to site directed mutagenesis so that codons 250, 254, 257, and 258 encode alanines rather than leucines, generating Swi6^1-273^ΔNES-GFP. Cells were synchronized as in Fig. 2 and observed by direct fluorescence (GFP) and phase-contrast (phase) microscopy at the time intervals indicated after release from a-factor arrest. (E) Nuclear export of Swi6^1-273^ΔNES-GFP is intermediate between Swi6^1-273^-GFP and Swi6^1-181^-GFP. Plasmids expressing Swi6^1-273^ΔNES-GFP, Swi6^1-273^-GFP, and Swi6^1-181^-GFP were expressed in synchronized cells and the percent of cells with nuclear GFP fluorescence was determined for each Swi6 fusion at 20-min intervals after release from a-factor. Error bars depict standard error of the mean from at least three experiments.

Proteins that are exported from the nucleus by exportin Crm1/Xpo1 typically contain a leucine-rich nuclear export signal (NES) or associate with another NES-containing polypeptide (Kirli et al., 2015; Stade et al., 1997; Thakar et al., 2013; Xu et al., 2010). Close inspection of the amino acid sequence found between Swi6 residues 181 and 273 revealed a leucine-rich region from amino acids 250 – 258 that aligns closely with NES sequences from other Crm1 export cargoes (Fig. 2C). In addition, the leucine-rich region spanning amino acids 250 – 258 is highly conserved across hemiascomycetes, with relatively rare amino acid changes within the putative NES relative to areas flanking the leucine-rich sequence (Fig. 2D).

To determine if this putative Crm1 NES is necessary for normal Swi6 localization throughout the cell cycle, we mutagenized Swi6^1-273^–GFP so that leucines at amino acids 250, 254, 257, and 258 were all altered to alanines (Swi6^1-273^ΔNES-GFP). Cells expressing Swi6^1-273^-GFP, Swi6^1-181^-GFP, and Swi6^1-273^ΔNES–GFP were then treated with a-factor and localization observed for synchronous populations of cells. Similar to the Swi6^1-181^–GFP truncation, Swi6^1-273^ΔNES–GFP remained predominantly nuclear as cells progressed from G1 into S-phase, although some cytoplasmic redistribution was observed (Fig. 2E). To quantify these changes, we determined the percentage of cells expressing primarily nuclear fluorescence at 20-min intervals after release from a-factor (Fig. 2F). Sixty min after a-factor release, fewer than 10% of cells expressing Swi6^1-273^-GFP exhibited nuclear fluorescence, compared to over 45% of cells expressing Swi6^1-181^-GFP, confirming the increase in nuclear retention of Swi6 lacking residues 181-273. Interestingly, cells expressing the Swi6^1-273^ΔNES-GFP exhibited an intermediate level of Swi6 export, with 26% of cells exhibiting primarily nuclear fluorescence 60 min after a-factor release. These data suggest that the leucine residues found within the predicted NES sequence in Swi6 are important but not essential for Swi6 nuclear export.

### Crm1 is an exportin for Swi6

The exportin Crm1/XpoI associates directly with leucine-rich NES sequences on cargo proteins in the presence of Ran-GTP in order to facilitate the translocation of cargoes out of the nucleus (Fornerod et al., 1997; Stade et al., 1997). If the leucine-rich domain found within Swi6 functions as an NES for Crm1, the Swi6 and Crm1 proteins should physically associate. To determine if there is a physical interaction between Crm1 and Swi6, we tested for protein-protein interactions *in vivo* using the yeast two-hybrid system (Fields & Song, 1989). Two-hybrid plasmids were generated that express full-length Swi6 in-frame with a *GAL4* transcription activation domain (Swi6-AD) and full-length Crm1 in-frame with a *GAL4* DNA binding domain (Crm1-BD). These plasmids were then introduced into cells either in combination or with an empty vector control containing only a *GAL4* DNA binding domain (pOBD-2) or transcription activation domain (pOAD; (Cagney et al., 2000)). Cells were then assayed for Crm1-Swi6 interactions using two different reporter genes (Fig. 3). We first used an *ADE2* two-hybrid reporter, which allows cells to grow on media lacking adenine if Swi6-AD and Crm1-BD interact *in vivo*. Cells expressing both Swi6-AD and Crm1-BD activated the *ADE2* reporter and exhibited robust growth on the –Ade selective media. In contrast, cells expressing any combination of either Swi6-AD or Crm1-BD with an empty vector did not grow on –Ade media. These data provide evidence that Swi6 and Crm1 physically interact *in vivo*. Interestingly, the co-expression of Swi6-AD and Crm1-BD reduced cell growth on non-selective media (Fig. 3A, left plate), while independent expression of either allowed for robust growth, providing additional genetic evidence for an interaction between the chimeric Swi6 and Crm1 proteins.

**Figure 3.**
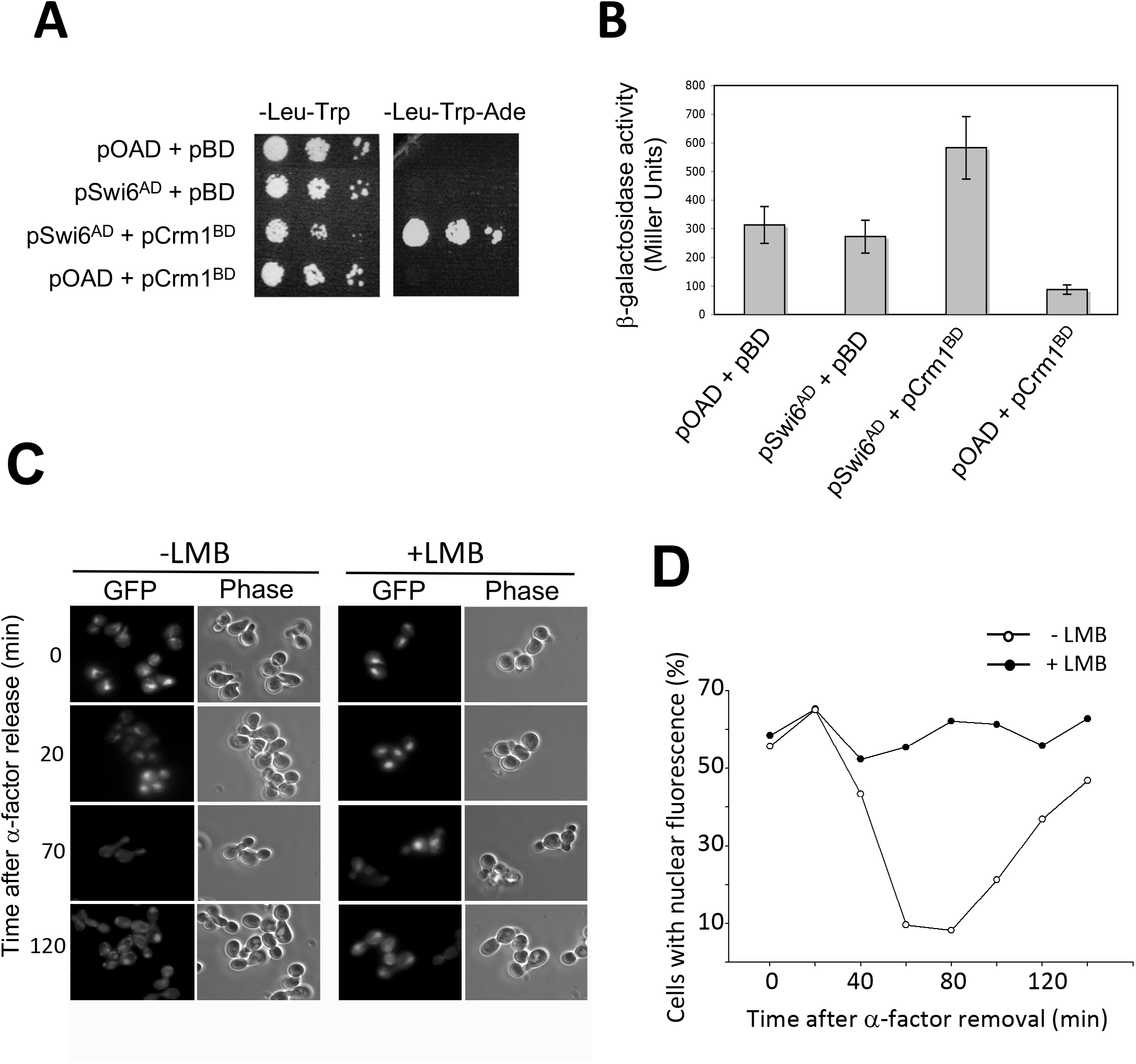
Swi6 associates with Crm1 and is requires Crm1 activity for efficient nuclear export. Plasmids containing Swi6 fused to a transcription activation domain (Swi6^AD^) and Crm1 fused to the LexA DNA-binding domain (Crm1^BD^) were co-transformed into two-hybrid reporter yeast strains together or in combination with two-hybrid vectors lacking the fusion (pAD or pOBD-2). (A) Transformed cells containing only empty vectors (pAD + pOBD-2), Swi6^AD^ and empty vector (pSwi6^AD^ + pOBD-2), Swi6^AD^ and Crm1^BD^ (pSwi6^AD^ + pCrm1^BD^), or empty vector plus Crm1^BD^ (pAD + pCrm1^BD^) were plated on media to select for transformation with both plasmids (Leu-Trp) and media to select for a two-hybrid interaction by complementation of an *ADE2* mutation (-Leu –Trp –Ade). Serial dilutions were plated on selective media and photographed after 72 hrs at 30°C. (B) Two-hybrid reporter yeast cells were transformed as above, grown in liquid media to select for both plasmids, lysed, and assayed for β-galactosidase activity as a marker for two-hybrid interaction. Data depict averages of three or more assays. Error bars represent standard error of the mean. (C) Yeast cells containing a mutation in Crm1 (Crm1^LMB^, (Neville & Rosbash, 1999)) that makes the protein susceptible to inhibition by leptomycin-B (LMB) were transformed with a plasmid expressing Swi6^1-273^-GFP. Overnight cultures in log-phase growth were arrested in G1 using a-factor and treated with LMB for 1 hr (+LMB) or left in media lacking drug (-LMB). Cells were then rinsed with +LMB or –LMB media lacking a-factor and observed by fluorescence microscopy. Cells in the absence (-LMB) or presence (+LMB) of leptomycin-B were photographed in a-factor (0 min) and 20 min, 70 min, and 120 min after release from cell cycle arrest. (D) Cells were treated as in (C) and the percent of cells with Swi6^1-273^-GFP nuclear fluorescence was determined at 20 min intervals after a-factor release.

To confirm this two-hybrid interaction between Swi6 and Crm1, we performed a quantitative assay for expression of an interaction-dependent LacZ reporter. Using the same combinations of two-hybrid plasmids described above, we lysed cells and assayed for LacZ expression using a colorimetric assay for β-galactosidase activity (Fig. 3B). Yeast cells containing both the Crm1-BD and Swi6-AD plasmids had a significantly higher expression of the LacZ reporter than cells containing the Crm1-BD and empty vector control (Tukey multiple comparison of means, p = 0.005). Crm1-BD/Swi6-AD co-expression also resulted in more than two times the β-galactosidase activity of cells expressing just Swi6-AD or empty vector, although variation between samples limited statistical significance (p = 0.1 and p = 0.07, respectively). These data provide further evidence that Swi6 and Crm1 physically associate, consistent with a role for Crm1 in Swi6 nuclear export.

To test if the exportin Crm1 is essential for Swi6 nuclear export, we introduced Swi6^1-273^-GFP into yeast cells expressing a leptomycin B (LMB)-sensitive form of Crm1 (Crm1^LMB^; (Neville & Rosbash, 1999)) as their only source of Crm1 protein. LMB associates directly with Crm1^LMB^, blocking Crm1 interaction with NESs and thus inhibiting nuclear export of proteins containing a Crm1 NES, but not affecting the nucleocytoplasmic shuttling of proteins transiting the NPC independently of Crm1 (Kudo et al., 1999; Neville & Rosbash, 1999). Crm1^LMB^ cells were treated with a-factor, incubated with LMB, then released from cell cycle arrest. Swi6^1-273^-GFP localization was observed over time by fluorescence microscopy (Figure 3C, D). While Swi6^1-273^-GFP, like full-length Swi6 (Sidorova et al., 1995), is normally nuclear during G1 and S-phases and cytoplasmic during G2 and M, Crm1^LMB^ cells treated with LMB retained Swi6^1-273^-GFP in the nucleus throughout the cell cycle. Thus, Crm1 activity is necessary for cell cycle-dependent changes in Swi6 localization and specifically for Swi6^1-273^-GFP nuclear export prior to G2-phase of the cell cycle. These data, along with evidence from Queralt and Igual (2003), provide evidence that Swi6 has two exportins, Crm1 and Msn5, that influence its coordinated, cell cycle-dependent export from the nucleus.

### Swi6 export by Crm1 is important for MBF activity

To determine if Swi6 export by each of the two exportins, Msn5 and Crm1, functions in distinct Swi6 activities, we investigated SBF and MBF activation in the absence of Crm1-mediated export. Nuclear export of Swi6 by Msn5 has been shown to be important for SBF activity (Queralt & Igual, 2003). Thus, we first sought to determine if Crm1-mediated export of Swi6 was also necessary for SBF activation. If Swi6 export by Crm1 is critical for SBF activity, then Swi6 harboring a Crm1-specific NES mutation should reduce expression of SBF targets. To determine if alteration of the Swi6 Crm1-NES sequence affects transcription activation by SBF, we altered a plasmid expressing wild type *SWI6* so that each leucine in the NES between residues 250 and 258 was converted to an alanine, making full-length Swi6ΔNES. We then expressed full length Swi6 and Swi6ΔNES in cells lacking endogenous *SWI6* and containing a chimeric reporter construct with an SBF-responsive promoter element upstream of a LacZ reporter (Costanzo et al., 2003).

Activation of gene expression for the SBF-LacZ reporter was subsequently assayed by measuring β-galactosidase activity in each strain (Fig. 4A). We observe that SBF activity is not significantly different between cells expressing *SWI6* and *swi6ΔNES* (p = 0.95), while a complete deletion of *SWI6* (*swi6Δ*) results in a statistically significant (p < 0.005) 10-fold reduction of SBF reporter activity.

**Figure 4.**
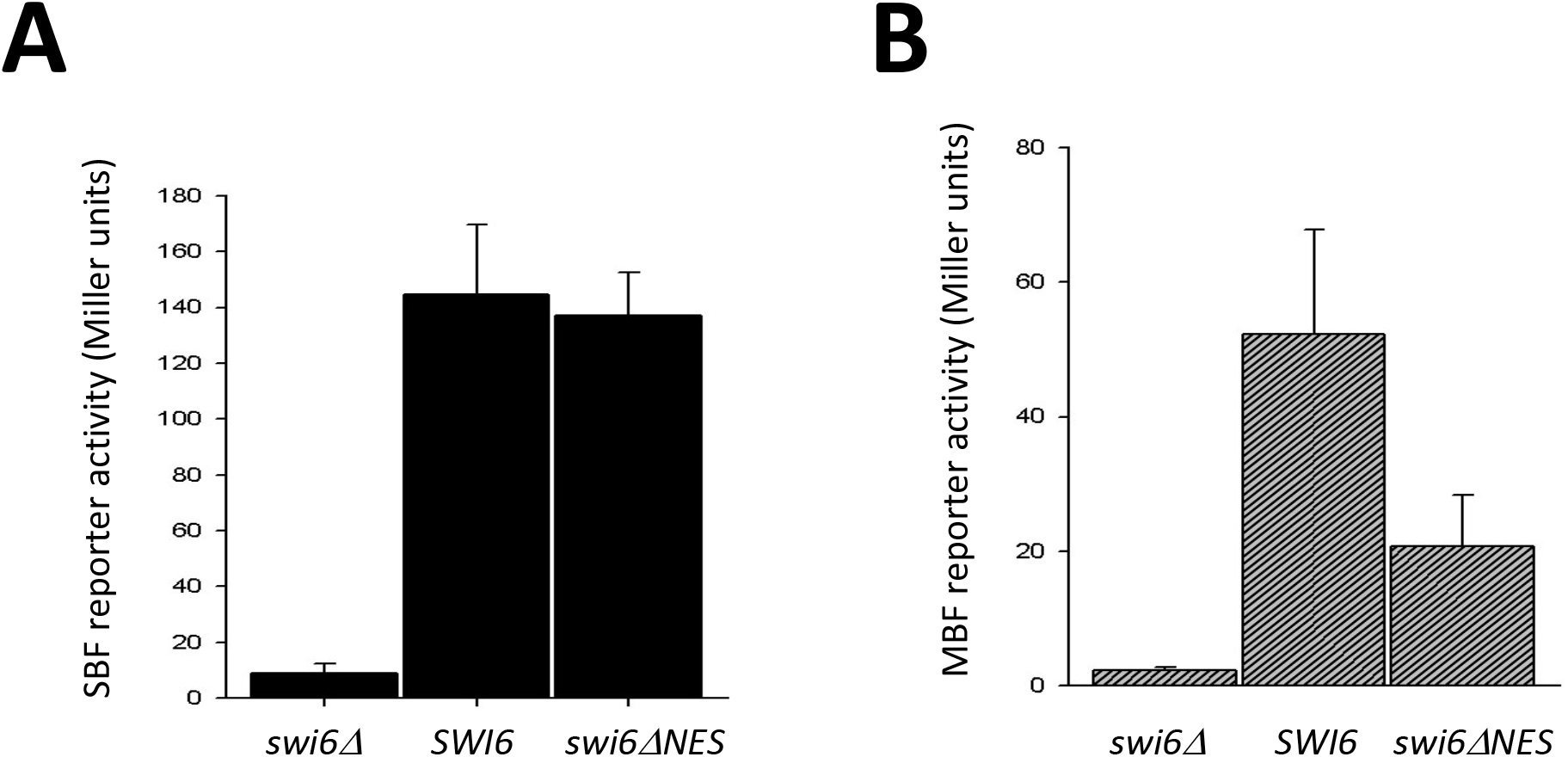
Alteration of the leucine-rich NES in Swi6 reduces MBF-mediated expression, but not SBF-mediated expression. Yeast containing a *swi6Δ* deletion were transformed with empty-vector (*swi6Δ*), a centromeric plasmid expressing wild-type *SWI6* (*SWI6*), or a centromeric plasmid expressing full-length *SWI6* with each of the leucines in the leucine-rich NES converted to an alanine (*swi6ΔNES*). Each strain was also transformed with a reporter plasmid containing either an SBF-response element (A) or MBF-response element (B) upstream of the *LacZ* gene. Cells were grown to log phase, lysed, and extracts analyzed for β-galactosidase activity. Bars represent average of at least three assays for each strain. Error bars show standard error of the mean for each.

Given that Swi6 participates in both SBF and MBF activity, we sought to determine whether the ANES mutation also affects trans-activation by MBF. To this end, we expressed *SWI6* and *swi6ΔNES* in cells containing an MBF-LacZ reporter and β-galactosidase activity was measured for each (Fig. 4B). MBF activity undergoes a decrease (2.5-fold) in cells expressing *swi6ΔNES* as their only source of Swi6 when compared to Swi6-expressing cells. However, MBF reporter activity remains higher in the presence of *swi6ΔNES* than in *swi6Δ* cells lacking Swi6 protein. While a pairwise comparison of the difference between MBF reporter expression in the *swi6ΔNES* strain and that in either *SWIΔ* or *swi6Δ* cells is not statistically significant to a p-value of less than 0.05, analysis by ANOVA indicates statistically significant overall variation in MBF expression among the three strains (p = 0.01). This trend suggests that the elimination of the Crm1-dependent NES from Swi6 reduces but does not eliminate the ability of Swi6 to contribute to MBF activity. These data suggest that Crm1-mediated export of Swi6 is important for the transcription activation function of MBF, but not of SBF, providing evidence that Crm1 and Msn5 are each important for unique aspects of Swi6 function.

### Decreased Swi6 export results in increased cell size

Yeast cells lacking Swi6 exhibit an increased cell size phenotype (Wijnen et al., 2002). To test whether the Crm1 NES is necessary for Swi6 function in regulating yeast cell size, we expressed the *swi6ΔNES* mutant in cells lacking endogenous *SWI6* and measured asynchronous cultures of log-phase cells for average cell diameter (Fig. 5). As controls, we expressed wild-type *SWI6* or transformed cells with an empty plasmid control vector. Expressing wild-type *SWI6* in *swi6Δ* deleted cells produced nearly normal-sized cells (7.69 μm average diameter; SE = 0.22 μm) compared to wild-type (7.03 μm; SE = 0.07 μm). *swi6Δ* cells harboring an empty vector, however, were significantly larger (11.82 μm; SE = 0.68 μm; p < 0.00001). Cells expressing *swi6ΔNES* as their only source of Swi6 protein displayed an intermediate average diameter (8.95 μm; SE = 0.19 μm) that is significantly larger than wild-type cells (average 1.92 μm larger; p < 0.01), but is significantly smaller than cells lacking *SWI6* (average 2.87 μm smaller; p < 0.00005). Thus, the loss of a Crm1-NES in Swi6 (*swi6ΔNES*) results in increased cell size but not to the extent observed in a *swi6Δ* mutant.

**Figure 5.**
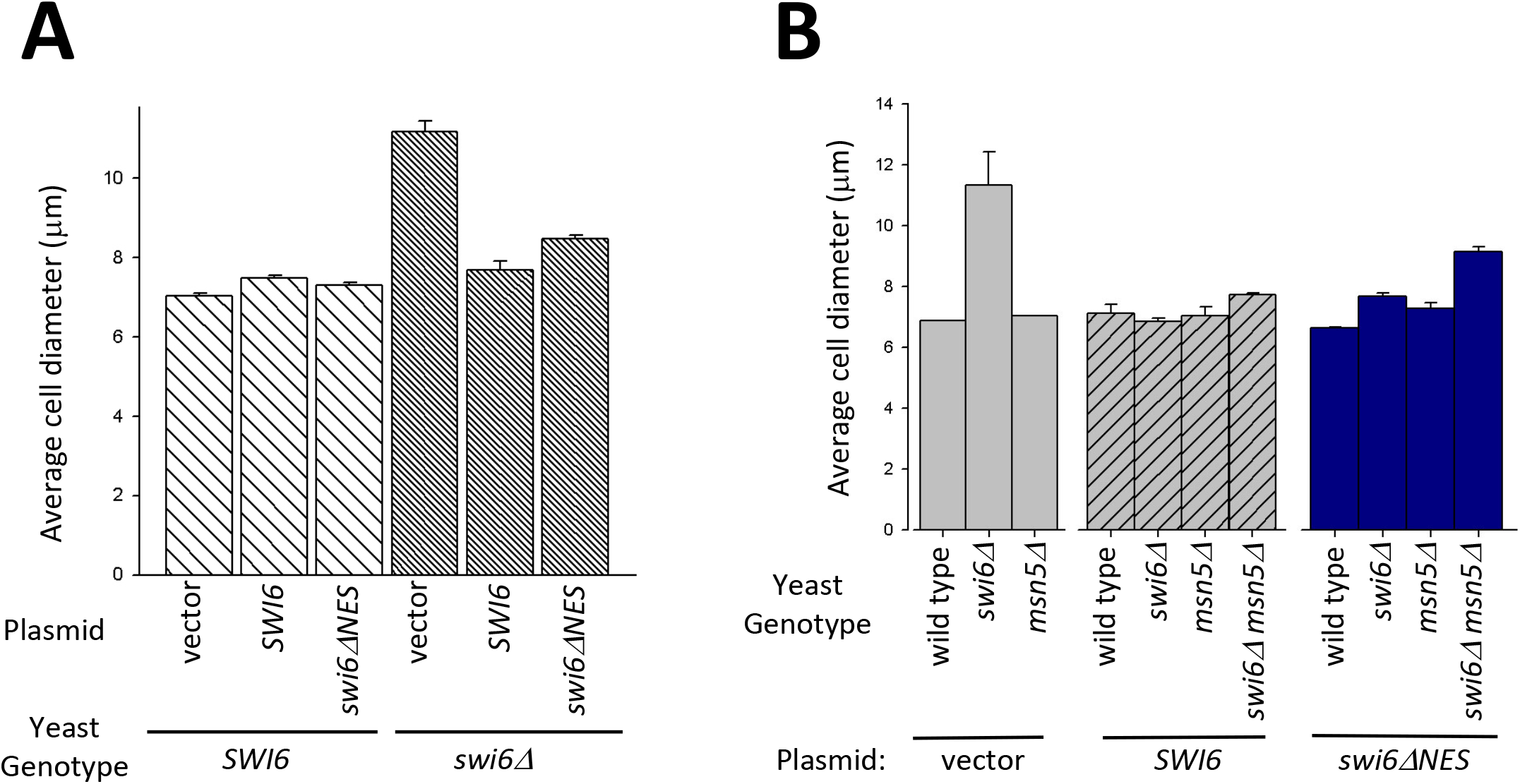
Swi6 nuclear export mutations result in increased cell size. (A) Cells lacking the Swi6 leucine-rich NES are larger than wild-type cells, but smaller than those completely lacking SWI6. Yeast cells containing genomic wild-type *SWI6* (*SWI6*; wide hashed bars) or a genomic deletion of *SWI6 (swi6Δ;* narrow hashed bars) were transformed with empty plasmid (vector), plasmid expressing wild-type *SWI6* (*SWI6*), or plasmid containing *SWI6* with a mutation that changes L -> A at amino acids 250, 254, 257, and 258 (*swi6ΔNES*). Bars represent the average cell diameter in μm. Error bars represent standard error of the mean. (B) Cells lacking both *MSN5* and the Swi6 NES are larger than either single mutant. Wild-type, *swi6Δ*, *msn5Δ*, and *swi6Δ msn5Δ* cells lacking a Swi6 plasmid (gray; *swi6Δ msn5Δ* not reported due to synthetic lethality), or containing a plasmid expressing *SWI6* (diagonally hashed) or *swi6ΔNES* (blue) were measured to determine average cell diameter. Error bars indicate standard error of the mean for each.

One explanation for the partial cell size phenotype observed in the *swi6ΔNES* mutant is that some Swi6ΔNES protein may be exported by Msn5/Kap142, which functions as an exportin for Swi6 (Queralt & Igual, 2003). Thus, we investigated whether a deletion of *MSN5* exacerbated the *swi6ΔNES* cell size phenotype. To this end, we generated *msn5Δ* yeast expressing *swi6ΔNES* in the absence of endogenous *SWI6*. Notably, while a *swi6Δ msn5Δ* double mutant is synthetically lethal (Queralt & Igual, 2003), such a double mutant expressing *swi6ΔNES* is viable, indicating that Swi6 lacking this NES sequence is competent to perform its essential activity. With respect to average cell diameter (Fig. 5B), a *swi6Δ* strain (11.35 μm; SE = 1.08 μm) is significantly larger than any other strain examined (Tukey multiple comparison of means; p < 0.0002), confirming previous observations that a complete loss of Swi6 function increases cell size (Wijnen et al., 2002). Interestingly, a *swi6Δ msn5Δ* strain expressing *swi6ΔNES* as its sole source of Swi6 is significantly larger (9.13 μm; SE = 0.18 μm) than either the *msn5Δ* (7.76 μm; SE = 0.034 μm; p < 0.0002) or *swi6ΔNES* (7.69 μm; SE = 0.092 μm; p < 0.006) mutant alone. That cells with deletions of both the *SWI6* NES and *MSN5* are significantly larger than wild type cells or either single mutant suggests that Msn5 is involved in mediating some Swi6 function independently of the Swi6 leucine-rich NES. In addition, *msn5Δ swi6ΔNES* double mutant cells are still significantly smaller than *swi6* null cells (p < 0.0002), indicating that Swi6 retains some activity that affects cell size in the absence of Msn5 or Crm1 export.

## Discussion

In this study, we analyzed the nuclear export of the Swi6 transcription factor in yeast and characterized how alterations in export affect phenotypes associated with Swi6 function. We observe that Swi6 shuttles between the nucleus and cytoplasm and that the exportin Crm1 bind to a leucine-rich NES-like sequence in Swi6 with similarity to canonical Crm1 NESs. Removal of the NES-like sequence from Swi6 alters the cell cycle dependent relocalization of Swi6 from the nucleus to cytoplasm, as does inhibition of Crm1 function using leptomycin B. Deletion of the Swi6 NES results in a reduction of MBF reporter activity, but not SBF activity, and results in cells that are significantly larger than wild type but not as large as those with a complete *swi6Δ* deletion. Taken together with the work of Queralt and Igual (2003), our studies show that Swi6 has at least two exportins, Msn5 and Crm1, that each associates with a distinct NES sequence, and that nuclear export by each impacts different aspects of Swi6 function.

Swi6 functions as a transcription factor and requires both nuclear import and export for its activity (Harreman et al., 2004; Kim et al., 2010; Queralt & Igual, 2003; Taberner & Igual, 2010). Here we provide evidence that individual Swi6 polypeptides are re-imported into the nucleus after export, suggesting that nucleocytoplasmic shuttling is a component of Swi6 function. Our heterokaryon-based shuttling data does not discern whether the nuclear export that occurs during this shuttling process is simply the result of passive diffusion from the nucleus or is actively mediated by exportin proteins. However, at 90kDa, Swi6 is above the 40-60kDs size threshold of the selectivity barrier for rapid, passive diffusion through NPCs (Li et al., 2016; Mohr et al., 2009; Timney et al., 2016). Thus, both the rapid rate at which nuclear accumulation in the second nucleus occurs and the additional evidence for a role for exportins in Swi6 function ((Queralt & Igual, 2003), this study) suggest that nuclear export of Swi6 is active, regulated, and an important process for its activity.

The work of Queralt and Igual (2003) linked the exportin Msn5 to Swi6 function. We have provided evidence that a second exportin, Crm1, is also important not only for Swi6 nuclear export, but also for its activity. Our data indicate that Swi6 and Crm1 interact in vivo and that this interaction takes place via a leucine-rich region within Swi6 with considerable similarity to canonical NESs recognized by Crm1. This observation is supported by high-throughput affinity data identifying Swi6 as being among those proteins interacting with Crm1 in a RanGTP-dependent manner (Kirli et al., 2015). Many nucleocytoplasmic transport cargoes associate with more than one karyopherin, including with different exportins functioning to translocate cargoes in the same direction across the NPC (Kirli et al., 2015; Mackmull et al., 2017). How might having more than one exportin mediating nuclear export be beneficial to a shuttling protein? The ability of a protein to be exported by multiple exportins may increase the steady-state rate of nuclear export, or may enhance the ability to conditionally regulate transport, either by regulating which exportin is present or which NES sequences are available to be bound to facilitate transport.

Alternatively, one or more exportins could be functioning largely to remove non-nuclear proteins from the nucleus to prevent aberrant nuclear function, as proposed for Crm1 (Kirli et al., 2015). In the case of Swi6, two different exportins, Msn5 and Crm1, associate with distinct NES sequences within the protein ((Kirli et al., 2015; Queralt & Igual, 2003), this study) but the function of these distinct interactions is unclear.

Swi6 is transported into the nucleus by more than one karyopherin, being imported both by importin a-β (Kap60/Srp1-Kap95) and by Kap120, each of which recognizes a distinct NLS sequence (Harreman et al., 2004; Kim et al., 2010; Taberner & Igual, 2010) and each of which appears to mediate nuclear import in response to a different stimulus (Geymonat et al., 2004; Harreman et al., 2004; Kim et al., 2010). Similarly, the import of STAT proteins in multicellular eukaryotes occurs via two different importins, each of which associates with a distinct NLS (Meyer et al., 2002). It is noteworthy that the type of karyopherin responsible for import determines which genes the STAT transcription factor will activate (Meyer et al., 2002), suggesting that importins can function in the regulation of interaction with downstream targets after translocation across the NPC.

Here we provide evidence that the nuclear export mediated by Crm1 and Msn5 may have distinct roles in Swi6 function. While a yeast mutant lacking Msn5 has a significantly reduced level of SBF-mediated, but not MBF-mediated, reporter gene expression (Queralt & Igual, 2003), yeast expressing Swi6 lacking the Crm1 NES as their only source of Swi6 exhibit significantly reduced expression of an MBF reporter, but no reduction of an SBF reporter (Fig. 6). Our data suggest that Crm1-mediated export of Swi6 is important for the transcription activation function of MBF, but not of SBF, providing evidence that Crm1 and Msn5 are each important for a different aspect of Swi6 function.

The observed changes in cell size provide further evidence that both Msn5 and Crm1 are important, but not essential, for Swi6 function. A deletion of *SWI6* results in a nearly 50% increase in average cell diameter in yeast ((Queralt & Igual, 2003; Wijnen et al., 2002); Fig. 7A and B). We observe here that independently eliminating Msn5-mediated export via an *msn5Δ* deletion or reducing Crm1-mediated Swi6 export through mutagenesis of the leucine-rich NES of Swi6 results in a significant increase in cell size over wild-type, but not nearly the increase observed in *swi6Δ* cells. This observation suggests that both exportins have some role in cell size regulation in yeast and, at least in the case of Crm1, that Swi6 is involved in mediating this role.

Importantly, we observe that a simultaneous reduction in both Msn5- and Crm1-mediated export of Swi6 results in a larger cell size than does reducing either individually (Fig. 7B), providing further evidence that both exportins contribute to Swi6 function. However, the *msn5Δ swi6ΔNES* cells are viable, unlike *msn5Δ swi6Δ* cells, and have an average diameter smaller than *swi6Δ* alone, indicating that Swi6 retains some function, even when both Msn5 and Crm1 transport are reduced. There are several possible explanations for this retention of Swi6 function in the *msn5Δ swi6ΔNES* double mutant. One possibility is that nuclear export may not be necessary for all functions of Swi6, even though Msn5- and Crm1-mediated export are each important for some independent transcriptional activities ((Queralt & Igual, 2003); Fig. 6). Alternatively, there may be some residual Swi6 export occurring in the *msn5Δ swi6ΔNES* double mutant, either via Crm1, some other exportin, or passive diffusion through the NPC. Our data indicate that removing the Crm1-NES does not eliminate Swi6 export completely (Fig. 3C), suggesting that either Crm1 or some other exportin remains able to transport swi6ΔNES efficiently enough to retain the activity necessary for viability in the absence of Msn5. An interesting question is whether or not there is something functionally distinct about the Swi6 protein exported by Msn5 and that exported by Crm1, and whether this difference is mediated via a physical alteration, either through a post-translational modification, dimerization (or multimerization) with another polypeptide, or some other form of chemical or physical regulation of Swi6. It also remains possible that Swi6 export is not required for all Swi6 activity and that some transcription activation or other activity that impacts cell size remains functional even in the absence of Msn5 and Crm1-mediated Swi6 export.

The intracellular localization of Swi6 is important for regulating its activity as a transcription factor in *Saccharomyces*. This study brings the number of karyopherins involved in regulating Swi6 function to four (Kap-α/β and Kap120 importins; Msn5 and Crm1 exportins) and highlights the importance of regulated nucleocytoplasmic transport on transcription factor function. The data reported here indicate that Msn5 and Crm1 are both important for Swi6 function and, interestingly, that each is important for activating different Swi6 targets. Future studies should provide insight into the complex interplay between these nuclear import/export mechanisms, how the interaction between transport substrates and each of their importins and exportins is initiated and regulated, and how this interaction ultimately affects the activity of proteins that undergo nucleocytoplasmic shuttling.

## Materials and Methods

### Strains, plasmids, and growth conditions

Yeast strains used in this study are listed in Table 1. Yeast strains were cultured at 30°C unless otherwise stated. Yeast cell culture and media preparation were performed essentially as in Sherman et al. (1991). Yeast transformations were performed as described (Gietz & Schiestl, 2007). Plasmids used in this study are listed in Table 2. *E. coli* cells were cultured at 37°C and plasmid isolation was carried out using a Qiagen Plasmid Mini Kit (Qiagen, Germantown, MD) according to manufacturer instructions. pKBB353 (pSwi6^1-181^-GFP) was generated by PCR amplification of the *SWI6* promoter and first 543 nucleotides of the *SWI6* coding region using upstream primer 5’- TACCGGGCCCCCCCTCGAGGTCGACGGTATCGATAAGCTTGATATCGAATTCTTGCAGGTCACTGTCTTCTGCTTCACTC-3’ and downstream primer 5’- CTAATTCAACAAGAATTGGGACAACTCCAGTGAAGAGTTCTTCTCCTTTGCTATTCGGAGTGGAGTCACTCTCAGC −3’, followed by homologous recombination into *SpeI* digested pLDB350. pKBB354 (Swi6^1-273^-GFP) was generated in the same way using PCR primers 5’- TACCGGGCCCCCCCTCGAGGTCGACGGTATCGATAAGCTTGATATCGAATTCTTGCAGGTCACTGTCTTCTGCTTCACTC −3’ and 5’- CTAATTCAACAAGAATTGGGACAACTCCAGTGAAGAGTTCTTCTCCTTTGCTGCTGTCATTATTAAGGGATGTAGGC-3’ to amplify the promoter and first 819 nucleotides of *SWI6*. Plasmids pKBB354, pAC1202, and pAC1571 all were mutagenized to change codons encoding leucines to those for alanines at residues 250, 251, 254, and 258 using Quik-Change Site Directed Mutagenesis (Agilent Technologies, Santa Clara, CA) and primer 5’- GTTGTAAATGATAATGAACAGAAGATGAAAGCAGAGGCATTCGCACAACGGGCGGCATTTCCAGAAATTCAAGAAATGCCTACATCCC-3’, resulting in *Swi6ΔNES* plasmids pKBB416, pKBB418, and pKBB441, respectively. Full-length *SWI6* was cloned into pLDB235 behind the *GAL1* promoter by PCR amplification of *SWI6* from genomic DNA using oligonucleotides 5’- ATTGTTAATATACCTCTATACTTTAACGTCAAGGAGAAAAAACTATAATGGCGTTGGAAGAAGTGGTACG-3’ and 5’- GGTATCGATAAGCTTGATATCGAATTCCTGCAGCCCGGGGGATCCACTAGTTCTTTAGCAGCCGGATCCT TTG-3’, followed by homologous recombination to make pKBB484. Two hybrid plasmid pKBB486 (Crm1-BD) was constructed by PCR amplification of full-length CRM1 using oligonucleotides 5’- TGAAGATACCCCACCAAACCCAAAAAAAGAGATCGAATTCCAGCTGACCACCATGGAAGGAATTTTGGATTTTTCTAACG-3’ and 5’- TTTTCAGTATCTACGATTCATAGATCTCTGCAGGTCGACGGATCCCCGGGAATTGCCATGTAATCATCAAGTTCGGAAGG-3’, transformation into yeast with linearized pOBD-2, and homologous recombination. Plasmid pKBB491 (Swi6-AD) was generated in the same way using pOAD and the PCR product resulting from amplification of SWI6 using primers 5’- GATGATGAAGATACCCCACCAAACCCAAAAAAAGAGATCGAATTCCAGCTGACCACCATGGCGTTGGAAGAAGTGGTACG-3’ and 5’- GGGTTTTTCAGTATCTACGATTCATAGATCTCTGCAGGTCGACGGATCCCCGGGAATTGCCATGCCCTCATGAAGCATGC-3’. All plasmids were subjected to DNA sequencing to confirm correct nucleotide sequence in all inserts.

**Table 1.**
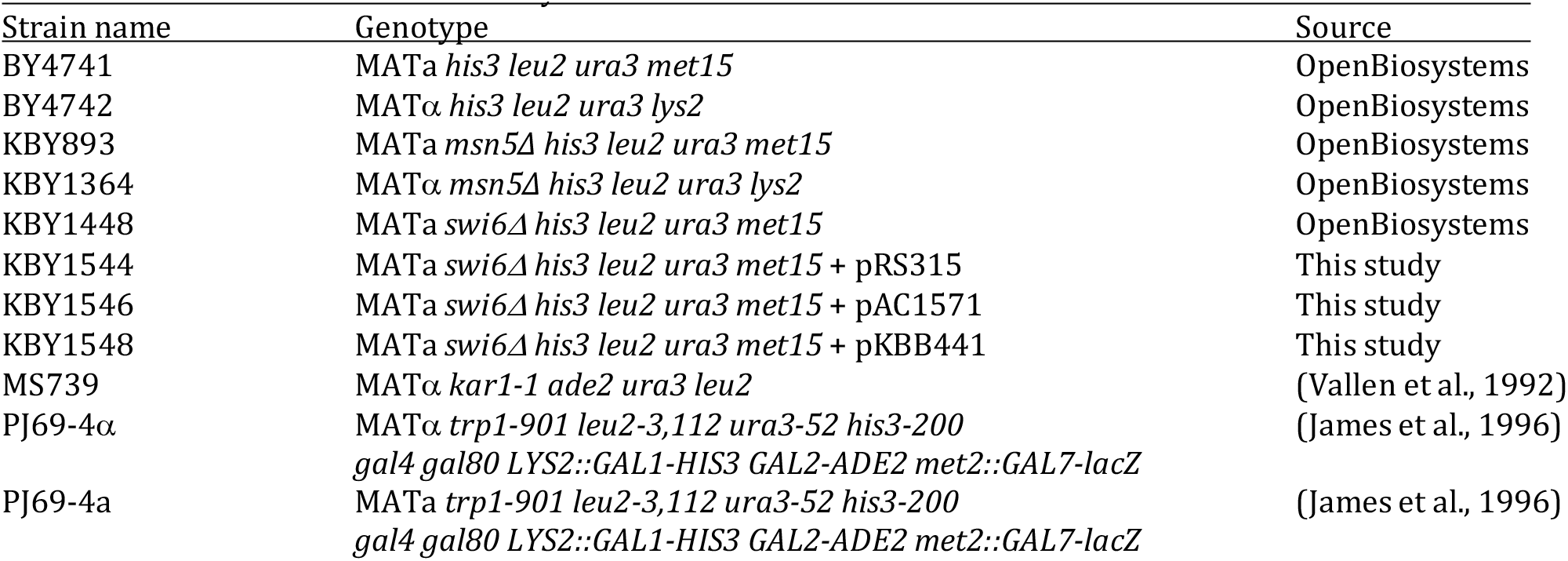
Yeast strains used in this study.

**Table 2.**
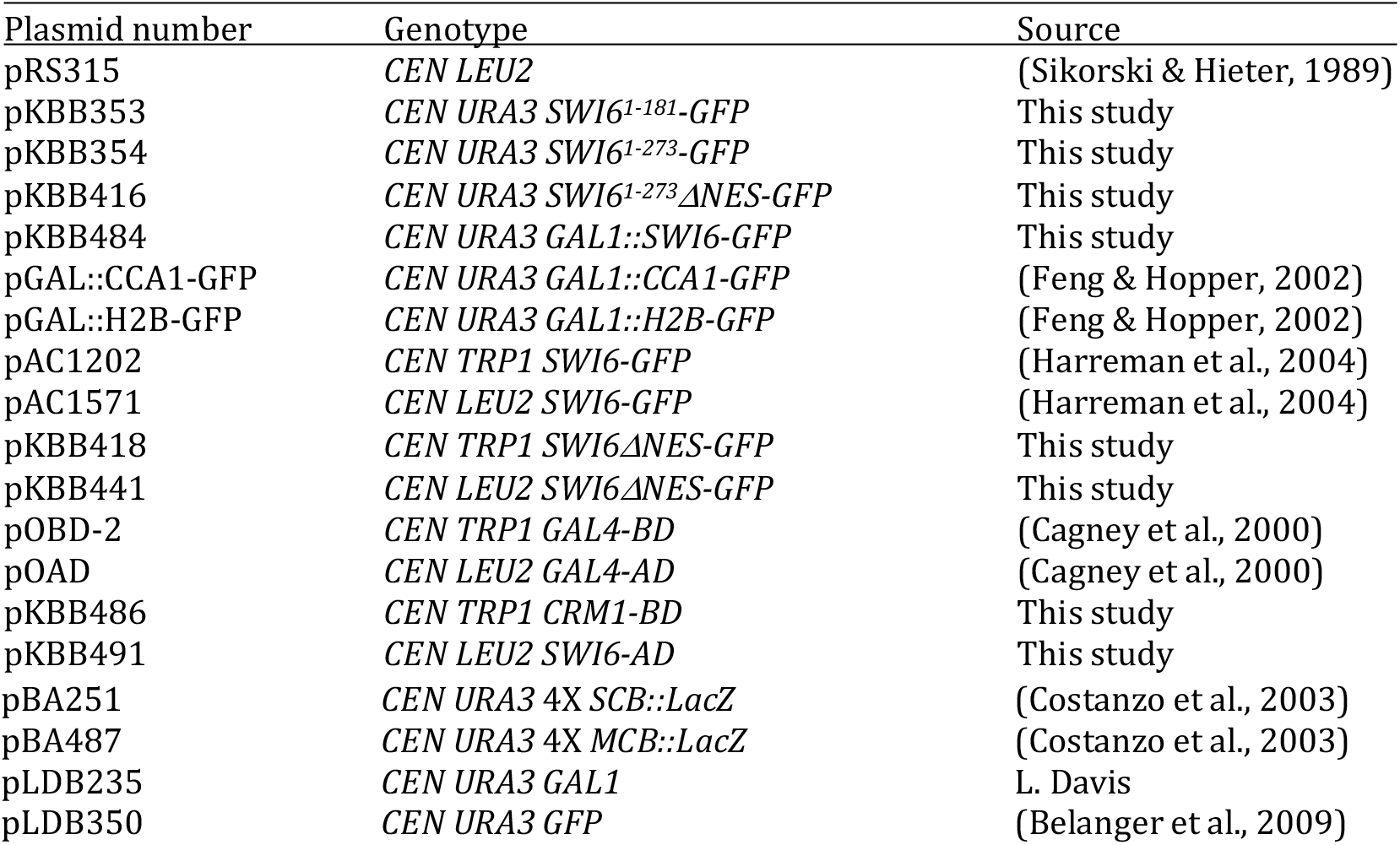
Plasmids used in this study

### Fluorescence microscopy

Fluorescence microscopy was used to visualize live cells containing each Swi6-GFP fusion. Cells were grown overnight at 30°C or 25°C in SD-Ura media. Two hours prior to microscopy, cells were diluted 20-fold. Cells were also examined after synchronization by a-factor treatment as described (Breeden, 1997). To release cells from arrest, a-factor was rinsed away twice with SD – Ura media, cells were incubated in selective media, and observation of nuclear versus cytoplasmic Swi6-GFP localization was recorded from more than 100 cells per timepoint. Nuclear fluorescence was defined as having fluorescence of greater intensity within the nucleus and a clearly defined transition between nucleus and cytoplasm. All cells without clearly defined nuclear fluorescence were scored as “cytoplasmic.” All assays were done in triplicate or greater. Photomicrographs were taken using Nikon E400 microscopes with SPOT RT_KE_ cameras and SPOT Advanced software (Diagnostic Instruments, Sterling Heights, MI).

### Heterokaryon assay

BY4741 wild type yeast containing either pKBB484 (*GAL1::SWI6-GFP*), pGAL::CCA1-GFP, or pGAL::H2B-GFP (Feng & Hopper, 2002; N. Mosammaparast et al., 2001) were grown overnight in SD –Ura media, then rinsed and resuspended in selective media with 2% raffinose for 3hrs at 30°C. After culture in raffinose, cells were washed twice with selective media containing 2% galactose and incubated 2 hrs at 30°C in SD–Ura with 2% galactose. Cells were then washed twice with SD-Ura with 2% glucose and incubated 2 hrs at 30°C. Heterokaryon formation and shuttling assays were performed essentially as described in Feng and Hopper (2002). Briefly, karyogamy mutant strain MS739 (*kar1-1*) and BY4741 cells containing pKBB484, pGAL::Cca1-GFP, or pGAL::H2B-GFP were vacuum filtered together onto a nitrocellulose filter and incubated on solid YPD media for 2 hrs to allow zygote formation. Cells were then scraped off of the filter, fixed in 70% ethanol, and DAPI stained for analysis by direct fluorescence microscopy and photomicroscopy as described above.

### Two-hybrid analysis

The yeast two hybrid system (Fields & Song, 1989) was used to test for protein-protein interactions. Yeast strains PJ69-4a and PJ69-4⍰ were respectively transformed with plasmids pOAD and pOBD-2; KBB491 (Swi6^AD^) and pOBD-2; KBB491 (Swi6^AD^) and KBB486 (Crm1^BD^); and pOAD & KBB486 (Crm1^BD^). The haploid transformants were mated and the resulting diploids were plated in 10-fold serial dilutions onto SD–Leu-Trp and SD–Leu-Trp-Ade plates and grown at 30°C. Cells were visually assayed for growth and photographed after 48 and 72 hr. Strains containing the same plasmid combinations were also assayed for β-galactosidase activity by cell lysis and colorimetric CPRG assay as described (Finn et al., 2013). Miller units of β-galactosidase activity were calculated using the method of (Miller, 1972). Statistical comparison of overall variation was performed by analysis of variance. Comparisons between conditions were performed by Tukey multiple comparisons of means at a 95% family-wise confidence level.

### SBF- and MBF-activity assays

Cells were assayed for SBF or MBF activity using the pBA251 (4X *SCB::LacZ*) and pBA482 (4X *MCB::LacZ*) plasmids (Costanzo et al., 2003). Briefly, yeast strains deleted for *SWI6* (*swi6Δ*) were co-transformed with either *SWI6* (pAC1571), *swi6ΔNES* (pKBB441), or empty vector (pRS315) plasmids in combination with pBA251 or pBA482. Transformants were isolated on SD – Leu –Ura selective media, cultured in liquid media, lysed, and assayed for *β*-galactosidase activity. Miller units of β-galactosidase activity were calculated as per (Miller, 1972). Statistical comparisons between strains within each group were performed using Tukey multiple comparisons of means at a 95% confidence level.

### Cell Size Analysis

To generate viable *swi6Δ msn5Δ* strains containing *swi6* mutations, we mated isogenic BY4741 and BY4742 yeast lacking *SWI6* and *MSN5*, respectively, and transformed the resulting diploid cells with empty vector or a plasmid expressing *SWI6* or *swi6ΔNES*. We then sporulated the transformed diploid cells and performed tetrad dissections on selective media to generate wild type, *swi6Δ, msn5Δ*, and *swi6Δ msn5Δ* cells containing each plasmid. Yeast strains were grown to A_600_ = 0.2 – 0.6 in 10ml of selective media at 25°C, and 50 μl samples were diluted in 150ml 1M NaCl and vortexed. Cell size was measured using an Elzone II Particle Size Analyzer (Micromeritics, Norcross, GA) as per manufacturer’s instructions. Three independent cultures of each yeast strain were measured and all cultures were analyzed in triplicate. Pairwise comparison of every combination of strains was made by Tukey multiple comparison of means at a 95% family-wise confidence level. Raw particle size data was analyzed using Microsoft Excel and SigmaPlot software.

## Acknowledgements

The authors wish to thank Anita Corbett, Anita Hopper, Laura Davis, Brenda Andrews, and members of Stan Fields’ lab for sharing of plasmids, strains, and reagents. We thank Kate Kokanovich and Amy Sullivan for technical contributions, Tim McCay and Larry Schweitzer for guidance and assistance with statistical analyses, undergraduate members of the Belanger lab for intellectual contributions and insightful discussions, and Bob Skibbens for critically reading the manuscript.

## Competing Interests

The authors declare no competing interests.

## Funding

This work was supported entirely by Colgate University through the Faculty Research Council, the Stuart Updike Endowed Undergraduate Research Fund, and generous funding in support of summer and academic year undergraduate research.

## Notes

### Competing Interest Statement

The authors have declared no competing interest.

